# Extensive Surveillance of Mosquitoes and molecular investigation of Arboviruses in central Iran; First Record of Molecular Identification of *Culex tarsalis* in Qom Province

**DOI:** 10.1101/2024.08.06.606915

**Authors:** Fatemeh Abedi-Astaneh, Hedaiatollah Raoofi Rad, Mohammad Hassan Pouriayevali, Tahmineh Jalali, Hassan Izanlou, Seied Abbas Hosseinalipour, Amir Hamta, Mohammad Es’haghieh, Mahdi Ebrahimi, Mohammad Ali Ansari-Cheshmeh, Mostafa Salehi-Vaziri, Asghar Talbalaghi, Ebrahim Abbasi

## Abstract

**Background:** Arboviruses are one of the greatest threats to animal and public health. *Culicidae* family is one of the most important vectors for the transmission of Arboviruses in the world. According to the geographical, demographic and climatic features of Qom city in Iran, it can be a suitable region for vectors and therefore transmission of Arboviruses.

**Methods:** In this study, which was conducted between 2019 and 2020 in different parts of Qom city, 83,414 mosquitoes were collected and after evaluating the species of mosquitoes based on morphological and molecular detection, the presence of alphaviruses, flaviviruses and phleboviruses were evaluated using genus specific RT-PCR assays.

**Results:** In this study, *Culex tarsalis, Culex theilerivoucher, Culex quinquefasciatus* and most importantly for the first time in Iran *Culex tarsalis* were detected. No alphavirus, flavivirus and phlebovirus infection was identified in collected mosquitoes.

**Conclusion:** Climatic and weather changes are the basis for the growth and spread of vectors and consequently the spread of arboviral diseases, and this issue seems to be very important on the necessity of increasing and continuing entomological and virological studies.

**Author Summary:** Arboviruses pose significant threats to animal and public health globally, with mosquitoes from the Culicidae family serving as key vectors in their transmission. This study focuses on Qom city, Iran, which, due to its unique geographical, demographic, and climatic characteristics, provides a potentially suitable environment for these vectors. Over the course of 2019 to 2020, researchers collected 83,414 mosquitoes across various locations within Qom. Through morphological and molecular analysis, they identified species such as *Culex tarsalis, Culex theileri*, and *Culex quinquefasciatus*, notably reporting *Culex tarsalis* in Iran for the first time. Using genus-specific RT-PCR assays, the study evaluated the presence of *alphaviruses, flaviviruses*, and *phleboviruses* in the collected samples but found no evidence of infection. These findings underscore the importance of ongoing entomological and virological research, particularly given the potential for climatic changes to influence vector growth and the spread of arboviral diseases.

## Introduction

Arthropod-Borne Viruses circulate between vertebrate hosts and arthropod vectors. Arboviruses has been considered as a serious public health challenge as they make up over than 17% of infections and have affected the lives of millions of people in a wide geographical area around the world(1). It is estimated that more than 520 viruses are transmitted by mosquitoes of which 130 are medically important(2, 3). The most important arboviruses mainly belong to the families of *Togaviridae* (genus Alphavirus), *Flaviridae* (genus Flavivirus), Orthobunyavirus, and Phlebovirus(4, 5). Mosquitoes such as *Culicidae* family are well-known vectors for arboviruses in worldwide(6). There are 3,100 species, 34 genera, and 3 subfamilies in the *Culicidae* family; The genuses/genera of the two subfamily Anophelinae (*Anopheles*) and Culicinae (*Aedes, Culex, Culiseta, Mansonia, Hemagogus, Sabethes* and *psoraphora*) which are the most important vectors of arboviruses(7).

In recent years, the risk of spreading mosquito vectors of arboviruses has increased significantly due to climate change and globalization, so that in the last decade, the world has seen widespread outbreaks and epidemics of these viruses, such as Chikungunya, West Nile, Zika and Dengue viruses(8). Apart from Went Nile virus, there have been no reports of autochthonous transmission of Zika, Dengue and Chikungunya viruses(9). However, imported cases of Chikungunya and Dengue viruses have documented from endemic countries mostly in East Asia(10, 11).

Owing to the fact that there is the close association between arboviruses and their mosquito vectors, the geographical distribution of vectors can be used to estimate the risk of emergence of arboviral infections and their outbreaks(12). Although several studies investigated the mosquitos’ fauna in Iran(13-15), there is no information regarding the arboviruses’ vectors fauna in Qom, Central Iran. The Qom is the main crossroad of Iran for transportation of goods and passengers of Iran. Qom is the pole of Shiism in the world, it has holy places such as the holy shrine of Bibi Fatima Masoumeh, the holy mosque of Jamkaran and the diversity of the presence of immigrants and pilgrims with ethnicities from most parts of Iran and nationalists from more than 100 countries. This city is the transportation crossroads of the country, so it is very important to control infectious diseases, carriers and disease reservoirs.

Since some Aedes are highly adaptable and adapt to different climatic, geographical, nutritional conditions, and have been able to spread in different tropical, subtropical and mild climate, and also because Aedes are urban insects and are moved by human vehicles, on the other hand, the vehicles of pilgrims and travelers from the southeastern neighbors of Iran sometimes arrive in Qom after traveling through the border provinces to Qom and sometimes directly of those countries; Therefore, it is possible to transfer Aedes mosquitoes of these vehicles to the city of Qom - The place of residence for a few days of the these people. The risk of spreading mosquito vectors of arboviruses has increased significantly due to climate change and globalization, so that in the last decade, the world has seen widespread outbreaks and epidemics of these viruses, such as Dengue, Zika, Chikungunya, and West Nile viruses. Because exact identification of mosquitoes is essential for controlling the species that are vectors of human and animal diseases(16). Therefore, this study was designed to discover the circulating mosquitoes in Qom province and investigate the alphaviruses, flaviviruses and phleboviruses infections in them.

## Materials and Methods

### Study Area

The Qom Province in Iran (Fig. 1) located in the centre of the country with an altitude of 900m above sea level. The population is approximately 1,365,000 and different types of climatic conditions are seen such as the mountainous in western and southern parts, semi-desert in central part and hot and dry desert in Eastern regions. The warmest months are June and July while January is the coldest month.

**Fig 1.**
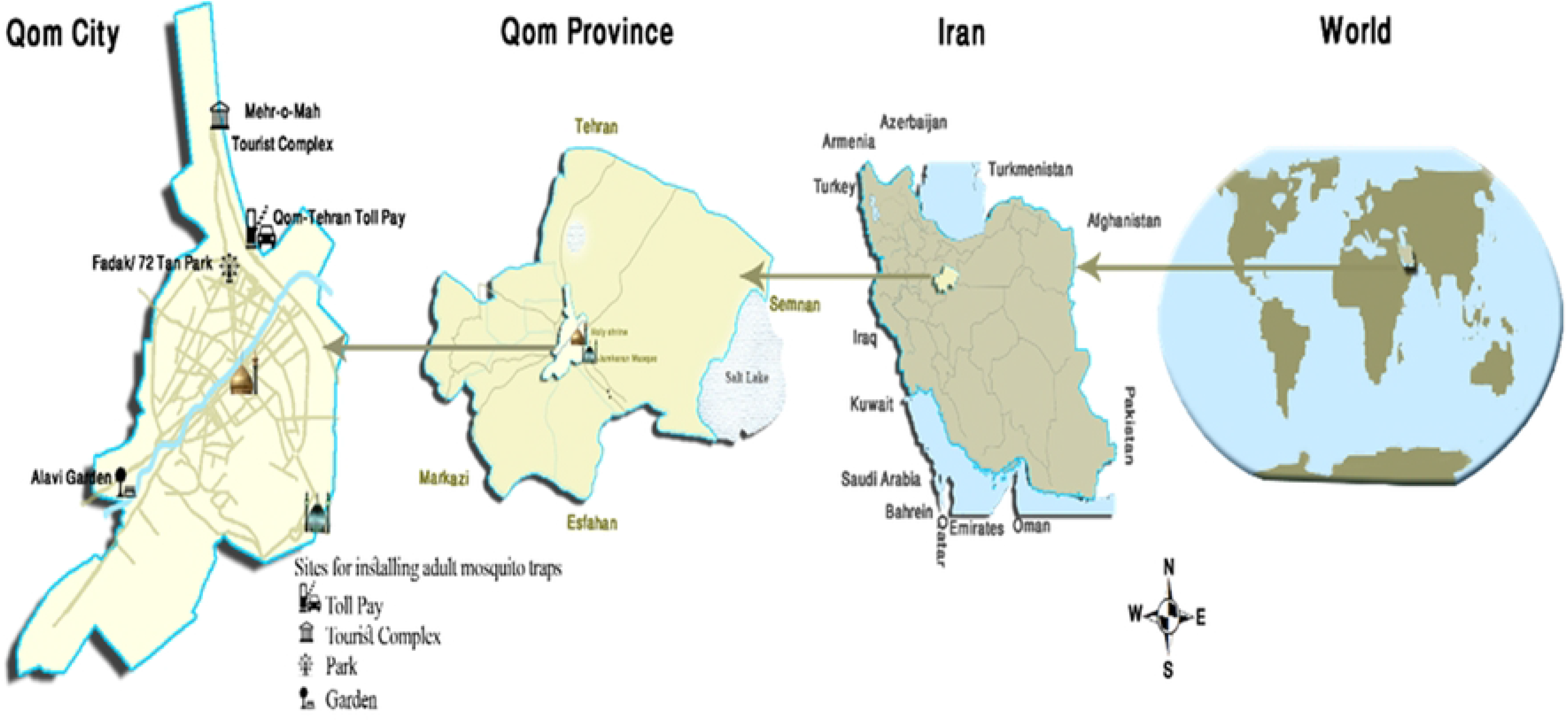
Qom Province in Iran, the study map.

#### Collection of Culicidae Mosquitoes

Eight traps (4 BG traps and 4 CO_2_ traps) were used to catch adult mosquitoes (Fig. 2). five separate ecological zones were identified for the field placement of 8 traps in the Qom province. The traps were placed at the entrances of the city of Qom, from the north, northwest, southeast and southwest of the country. one CO_2_ trap and one BG trap were placed in the Mehr-o-Mah rest area and tourist complex, one CO_2_ trap was placed in the Qom-Tehran toll plaza, one CO_2_ trap and one BG trap were placed in the 72 Tan Park, one BG trap were placed in the Maral Setareh tourist complex and one CO_2_ trap and one BG trap were placed in the Alavi Garden (Fig. 3).

**Fig 2.**
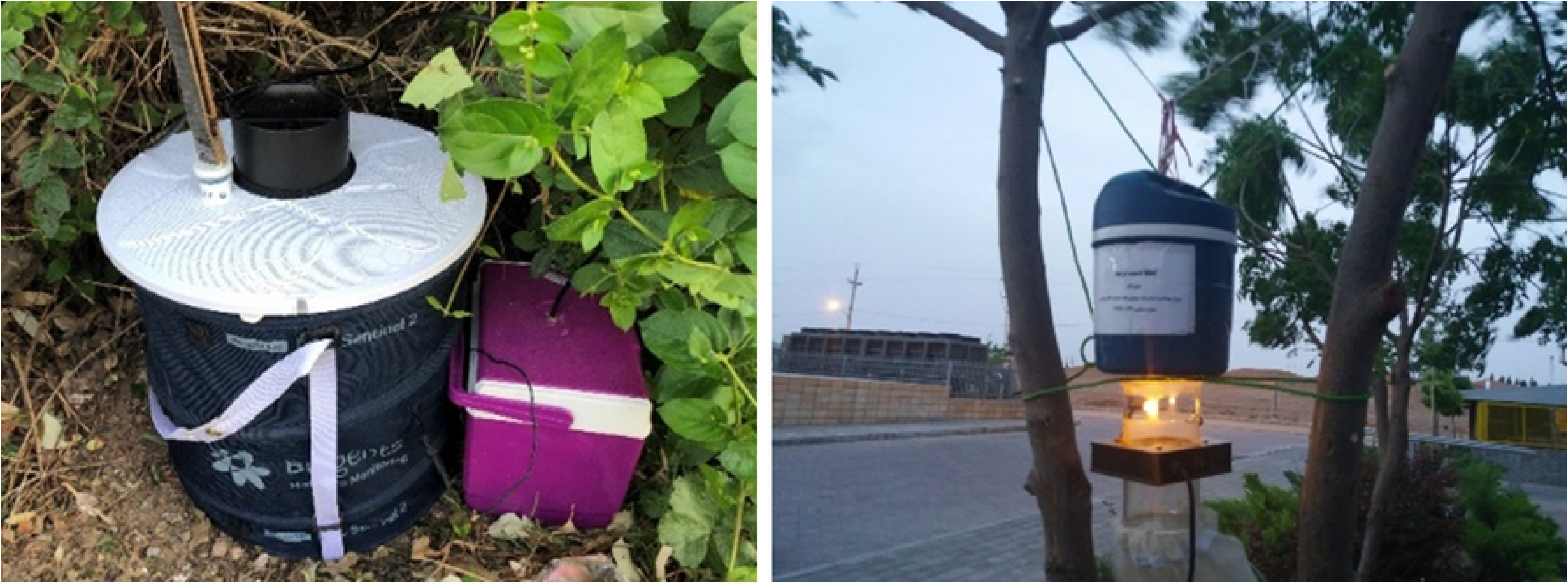
Left: BG trap and Right: CO_2_ trap.

**Fig 3.**
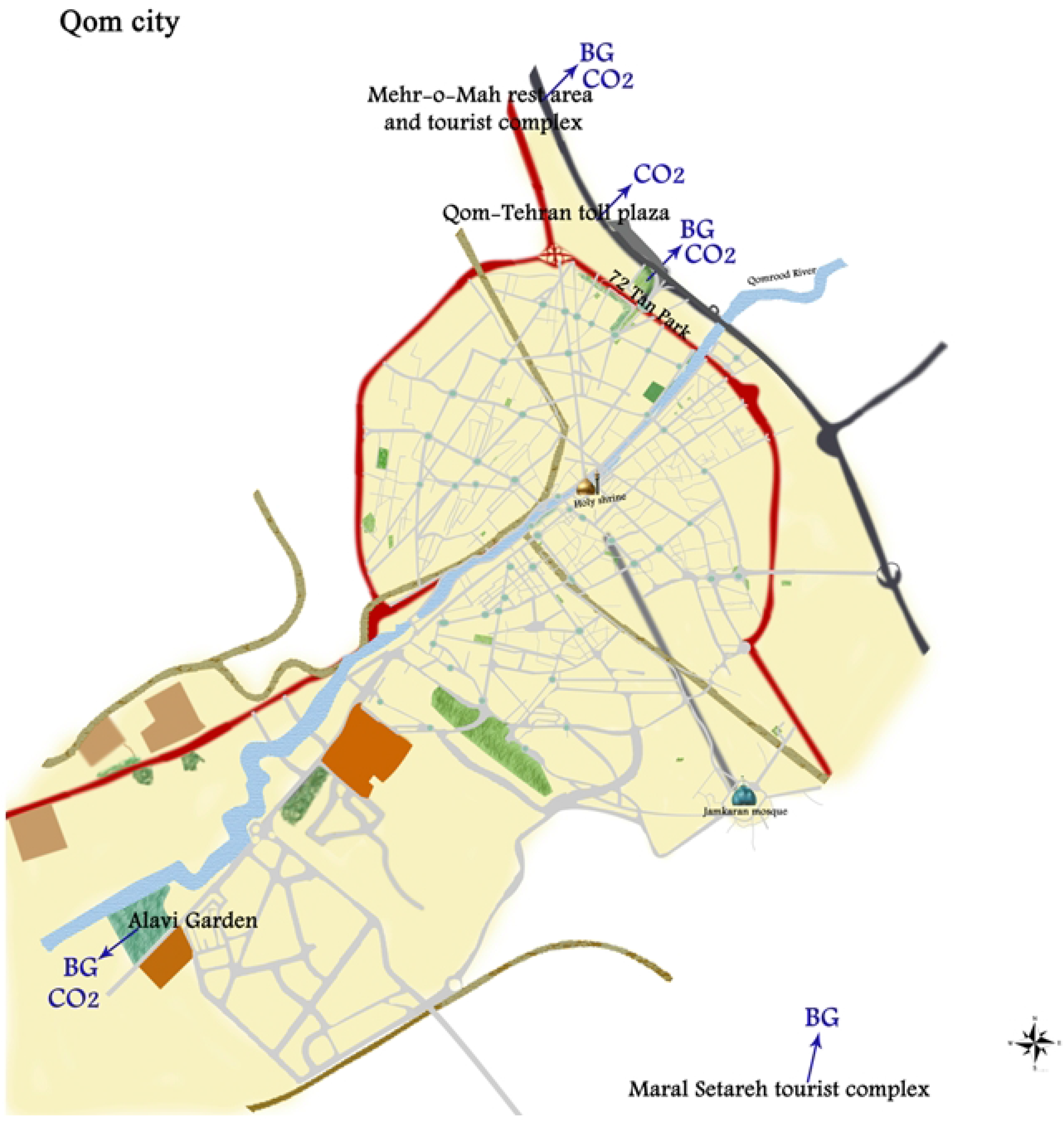
Map of the study area, Qom city, Qom Province, Iran, 2018.

The BG Traps were installed before sunrise. The CO_2_ traps installed at dusk and dismounted half hour after sunrise of the following day. In CO_2_ traps, light was used as an additional adult mosquito’s attractant. Traps were installed in public places, such as parks and tourist complexes. The *Culicidae* mosquitoes were morphologically identified within 1-2 hours and some of them were selected as pool and kept at liquid nitrogen for further investigation for Species of mosquitoes and possible presence of arboviral infection.

### Morphological Identification of Mosquitoes

Collected mosquitoes were transferred to the entomology laboratory placed in dry ice for a few minutes and microscopic diagnosis was done within 1-2 hours as previously described(17, 18). In microscopic diagnosis, diagnostic Keys and a 60X stereomicroscope lens were used. Undetectable or suspected mosquitoes and a number of identified mosquitoes were selected for molecular testing. Each pool contained an average of 40 mosquitoes of one species; of some species, only one mosquito was caught.

### Molecular identification of Mosquitoes

A Total of 20 pools of mosquitoes were subjected to molecular identification as well as morphological identification. These 20 pools were selected randomly and were sent to the department of Arboviruses and Viral Hemorrhagic Fevers (national ref lab), Pasteur Institute of Iran, with maintaining cold chain using dry ice and cold box for molecular investigations. All samples were stored at −70 ^°^C immediately after receiving for further analyses.

For molecular characterization of mosquitoes, PCR amplification and sequence analysis of 735 base pair fragment including Mitochondrial COI gene was carried out using SP ID F/SP ID R primers(19). Additionally, the specific primers for actin-1 (ActF and R)(20) were used. Briefly, mosquitoes were homogenized using Tissuelyser II (QIAGEN, Germany). The genomic DNA was extracted from pools using High Pure Viral Nucleic Acid Kit (Roche, Germany) according to manufacturer’s instruction. PCR was performed by the Hot StarTaq Master Mix Kit (Qiagen, Germany).

The final volume was 20 μL for each reaction performing one cycle at 95°C for 15 min, five cycles of 94°C for 40 sec, 45°C for 60 sec and 72°C for 60 sec and 35 cycles at 94°C for 40 sec, 51°C for 60 sec and 72°C for 60 sec and final extension 72°C for 10 min. PCR products (735bp for SP ID primers and 350bp for ACT primers) were sequenced bidirectional using applied biosystem 3130 Genetic Analyzer (Thermofisher, USA). The sequencing raw data was analysed by CLC Main Workbench software (CLC bio, Denmark) and using BLAST https://blast.ncbi.nlm.nih.gov/Blast.cgi) for confirmation, then all sequences data submitted to GenBank (https://www.ncbi.nlm.nih.gov/genbank) database (Table 1).

**Table 1.** Evaluation of mosquito detection by microscopic method and confirmation by sanger sequencing.

| NO. | Possible microscopic detection of species | The list of mosquitoes after sequencing and blasting | Accession number | Results |
| --- | --- | --- | --- | --- |
| 1 | *Culex pipiens* | *Culex pipiens* | OM714817 | Compatible |
| 2 | *Culex pipiens* | *Culex pipiens* | OM628998 | Compatible |
| 3 | ***Culex spp*** | ***Culex tarsalis*** | ON420215 | **Incompatible** |
| 4 | *Culex theileri* | *Culex theiler* | OM638648 | Compatible |
| 5 | *Culex modestus* | *Culex modestus* | OM638670 | Compatible |
| 6 | *Culex subochrea* | *Culex subochrea* | Not sequenced | - |
| 7 | *Culex perexiguus* | *Culex perexiguus* | Not sequenced | - |
| 8 | *Anopheles* | *Anopheles claviger* | OM638671 | Compatible |
| 9 | *Aedes caspius* | *Ochlerotatus caspius* | OM638672 | Compatible |
| 10 | *Aedes caspius* | *Ochlerotatus caspius* | OM638673 | Compatible |
| 11 | *Aedes caspius* | *Ochlerotatus caspius* | OM638674 | Compatible |
| 12 | *Aedes caspius* | *Ochlerotatus caspius* | Not sequenced | - |
| 13 | *Culex spp* | *Culex theilerivoucher* | OM638676 | **Incompatible** |
| 14 | *Culiseta longiareolata* | *Culiseta longiareolata* | OM638715 | Compatible |
| 15 | *Culex quinquefasciatus* | *Culex quinquefasciatus* | OM638717 | Compatible |
| 16 | *Culex spp* | *Culex quinquefasciatus* | OM714908 | **Incompatible** |
| 17 | *Culex quinquefasciatus* | *Culex quinquefasciatus* | OM714856 | Compatible |
| 18 | *Culex quinquefasciatus* | *Culex quinquefasciatus* | OM714858 | Compatible |
| 19 | *Culex quinquefasciatus* | *Culex quinquefasciatus* | OM714899 | Compatible |
| 20 | *Culex quinquefasciatus* | *Culex quinquefasciatus* | Not sequenced | - |

### Detection of Arboviruses

Homogenized mosquitoes were used for total RNA extraction using High pure viral RNA kit (Roche, Germany). The presence of Alphaviruses, Flaviviruses and Phleboviruses RNAs were investigated by Pan Phlebo RT-PCR using ppLF/ppL1R/ppL2R primes; Pan Flavi RT-PCR using MAMD/cFD2 primers and Pan Alpha RT-PCR usnig E1-S/E1-C primers(21). All RT-PCR reactions were done using Onestep RT-PCR kit (Qiagene, Germany). The final volume for each reaction was 25 μL performing one cycle at 50°C for 30 min, 95°C for 15 min, 40 cycles at 94°C for 30 sec, 53°C for 30 sec and 72°C for 45 sec and final extension 72°C for 5 min.

## Results

In this study, during 2019-2020, a total of 83,414 mosquitoes were caught. According to the morphological key, all categorized as *Culicidae* addition 933 Sandfly were caught. 27,057 *Culicidae* and 116 Sandfly were selected and kept at liquid nitrogen in 367 Pool including 355 *Culicidae* mosquitoes and 12 were *phlebotomine*. Since the microscopic and molecular diagnosis of *phlebotomine* were the same and no arboviruses were discovered in them, in this article, we will not continue the discussion about phlebotomines and about *Culicidae*, each pool contained an average of 76 mosquitoes of one species. To evaluate the results of morphological identification, 20 pools were subjected to morphological identification, as well.

The results of morphological and molecular characterization of mosquitoes were incompatible for three pools (Table 1). The pools no.3 and no.13 and no.16 were categorized as just Culex because were not recognise its species. In morphological identification, while the pools no.3 was characterized as *Culex tarsalis* after genomic analysis. According to the molecular assay, the pool no.13 was *Culex theilerivoucher*, and the pool no.16 was categorized as *Culex quinquefasciatus* in molecular assay.

In comparison, in microscopic method, of these 20 Pool, 3 Pool were misdiagnosed or incorrect. *Culex tarsalis* has been reported for the first time in Iran and the main reason for the lack of detection by the entomologist was the absence of this mosquito in the available diagnostic keys in Iran. Therefore, the Entomologist could only determine the genus (Fig. 3). The *Culex theileri* has a bright transverse band on the abdomen(22, 23), but in our study it was a rectangular band in the third abdominal band (Fig. 4&5) (Table 1). No Alphavirus, Flavivirus and Phlebovirus infection was identified in the collected mosquitoes.

**Fig 4.**
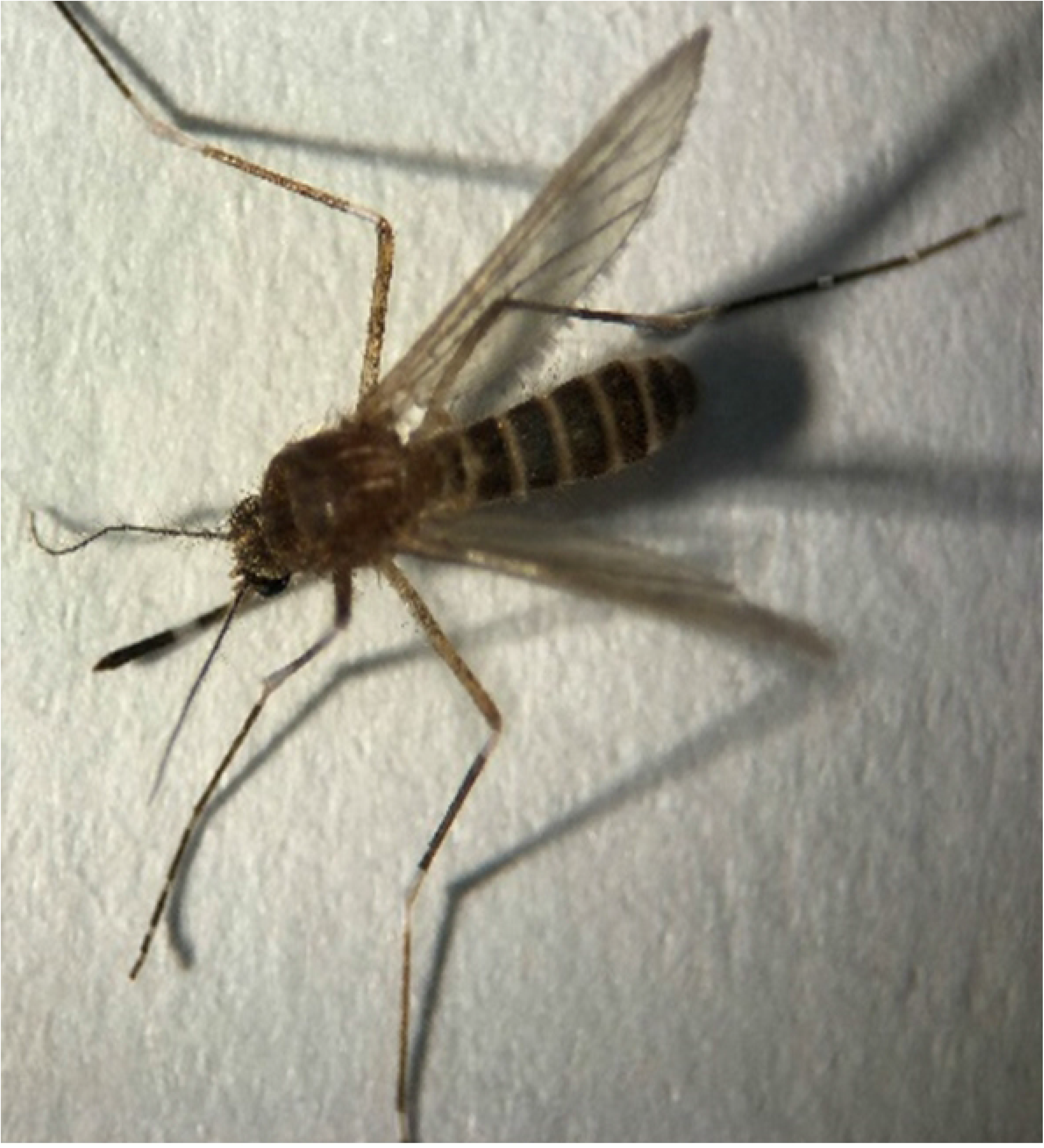
*Culex tarsalis. Cx. tarsalis* was not in the diagnostic keys of Iran, the Entomologist determined the genus and the national reference laboratory determine the species.

**Fig 5.**
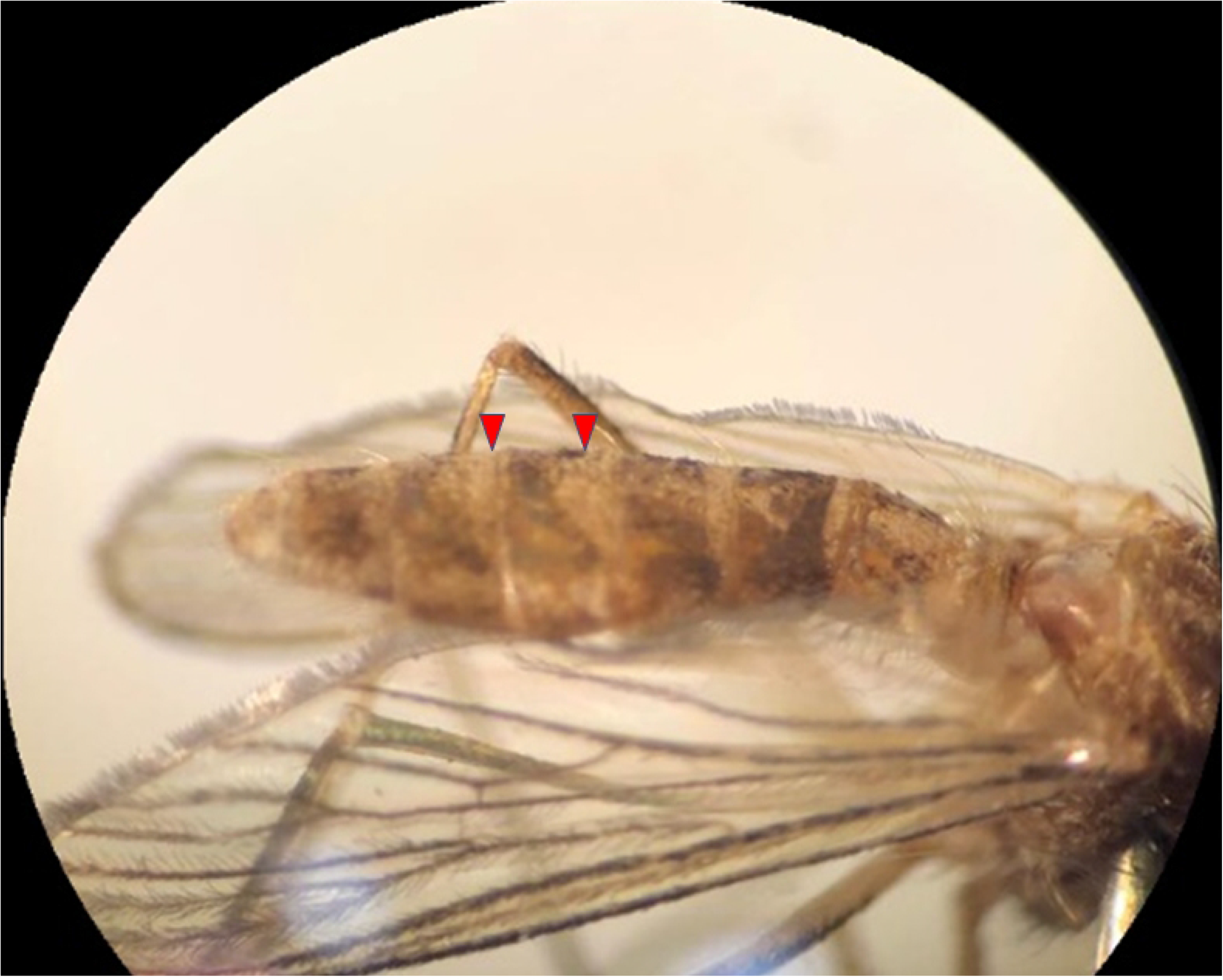
*Cx. theileri* (Unknown to Entomologist).

**Fig 6.**
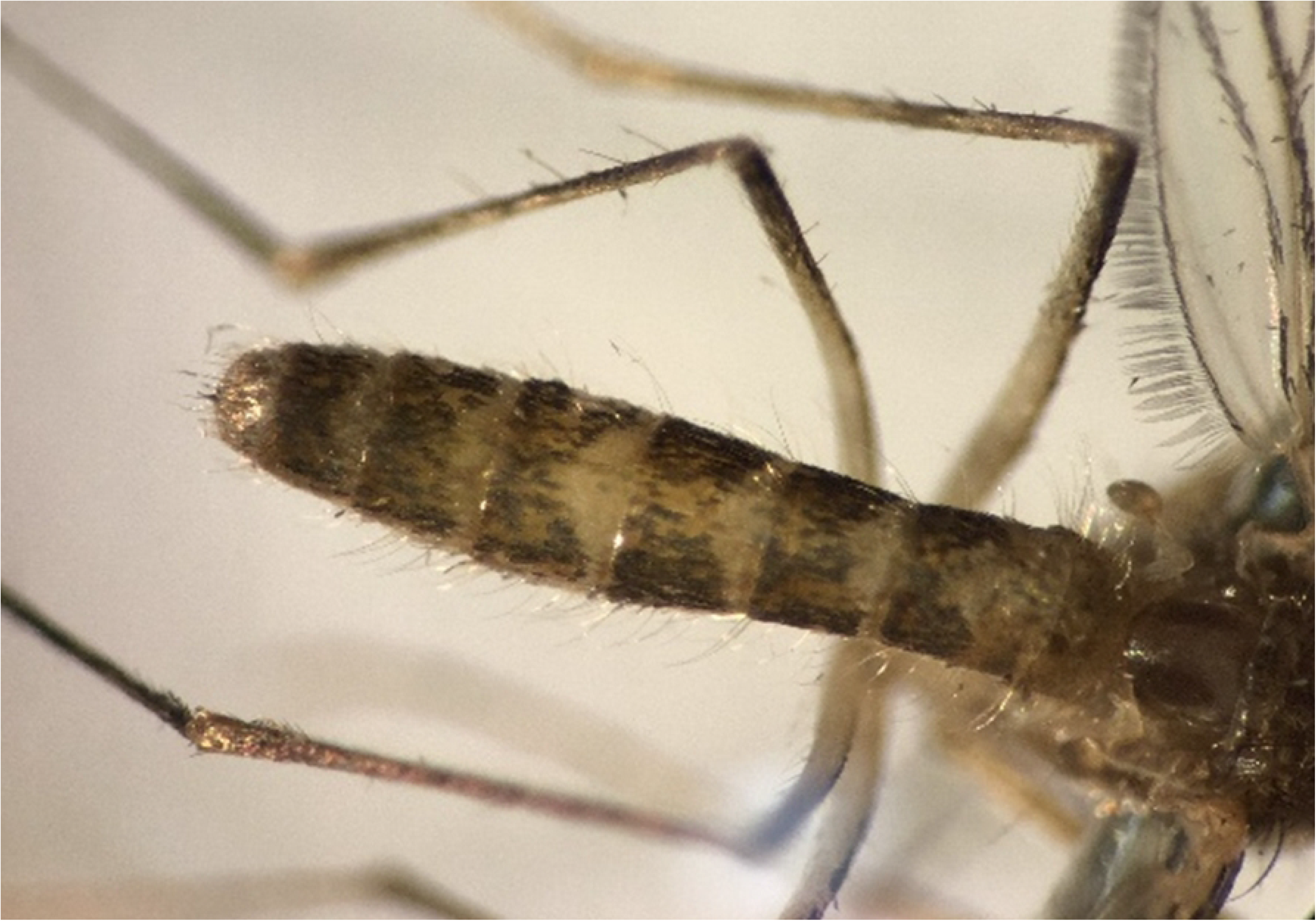
*Cx. theileri*

The *Culex theileri* has a bright transverse band on the abdomen. There is a rectangular band in the third abdominal band! Of course, it has the vertex of a bright triangle.

## Discussion

Considering that Qom province is a central city for the transit of goods to the south and north of the Iran, mosquitoes can be moved mechanically into or out of this region. In addition, a large number of tourists from Iran or other Muslim countries travel to this city for religious purposes and can be bitten by mosquitoes in those periods of residency.

In our study, 11 species were collected and identified which included *Culex quinquefasciatus, Culex pipiens, Culex theileri, Culex subochrea, Aedes (Ochlerotatus) caspius, Culex modestus, Culex theileri voucher, Anopheles claviger, Culex perexiguus, Culiseta longiareolata* and *Culex tarsalis* in Qom province. *Culex tarsalis* was identified for the first time from Iran. Asgarian et al. in fauna and larval habitat, from April to December 2019, identified three genera *Anopheles, Culiseta* and *Culex* in Kashan and also *Culex pipiens* (37.36%) and *Cx. Theileri* (26.10%) had highest abundance species(13). Nikookar et al. reported *Culex pipiens* as the most abundant species in fauna and larval habitat in Neka County, Northern Iran(24).

*Moradi-asl* et *al*, identified 13 species of mosquitoes in the form of four genus (Anopheles, Aedes, Culex and Culiseta) in north-western province of Iran and in this study, the Culex and *Culex Pippins* were the most abundant genus and species respectively(25).

In the current study, the most identified species is *Culex quinquefasciatus*, but Saghafipour et al. in Qom province reported four genera among Diptera: *Culicidae* which included 14 species and from them, *anopheles claviger* was most common species(26). Therefore, it seems that identification of different rates among the species can be due to different conditions and climatic variables. On the other hand, morphological studies alone are not sufficient to differentiate species and molecular methods can be helpful in identifying the species with high sensitivity. *Culex tarsalis* are strong flyers, covering distances of up to 4 km for blood meal and feeds from birds, cattle, horses and human(27).

Finding a new species of mosquito for the first time in Iran also warns that due to global warming and climate change, we must always wait for new species of mosquitoes as well as their establishment and so spreading emerging and re-emerging diseases. In addition, by their genetic and behavioural changes in each species, physical diagnosis become more difficult, and therefore molecular detection should always be used beside physical and morphological diagnosis.

Except for some molecular and serological reports for West Nile virus in Iran, Chikungunya virus and Dengue viruses have been reported as imported cases into Iran. In the presence study no Alphavirus, Flavivirus and Phlebovirus been reported(10, 19, 28-30).

However, due to climate change, there is a possibility that these viruses will spread in the future. Therefore, continuous entomological and virological studies beside to field studies on vectors in the region seem to be necessary.

## Conclusion

Due to the weather and climate changes that have occurred in Iran, it has caused the appearance of new species and even changes in the lifestyle of mosquitoes. Therefore, continuous studies in different geographical areas on viral vectors provide new information about the spread of arboviral diseases and their vectors.

## Acknowledgements

The authors would like to thank all managers and personnel of Qom Health Deputy, Qom University of Medical Sciences, the department of Arboviruses and Viral Hemorrhagic Fevers, Pasteur Institute of Iran; and the municipalities of Qom, Qom Green Space Organization, General Directorate of Cultural Heritage, Handicrafts and Tourism of Qom.

## Ethical considerations

This project was submitted in Qom University of Medical Sciences with code number 97959.

